# Comparison of Multiple Carbapenemase Tests Based on an Unbiased Colony Selection Method

**DOI:** 10.1101/2023.12.19.572349

**Authors:** Hsin-Yao Wang, Yi-Ju Tseng, Wan-Ying Lin, Yu-Chiang Wang, Ting-Wei Lin, Jen-Fu Hsu, Marie Yung-Chen Wu, Chiu-Hsiang Wu, Sriram Kalpana, Jang-Jih Lu

## Abstract

Carbapenemase-producing organisms (CPOs) present a major threat to public health, demanding precise diagnostic techniques their detection. Discrepancies among CPO tests have raised concerns, partly due to limitations in detecting bacterial diversity within host/specimen. We explored the impact of unbiased colony selection on carbapenemase testing and assessed its relevance on various tests. Based on “FirstAll” for unbiased colony selection to reduce bias, we compared modified carbapenem inactivation method/EDTA-modified carbapenem inactivation method (mCIM/eCIM), Carba5, the CPO panel, and multiplex PCR (M-PCR). Initially, we compared FirstAll to conventional colony selection for mCIM. Second, we used M-PCR as a reference, to evaluate test performance across seven CPO species. The results revealed that FirstAll selection improved carbapenemase detection, revising false-negative in 10.5% of *K. pneumoniae* isolates. In addition, 12.4% of CPOs tested positive for multiple carbapenemase genes. Both the Carba5 test and CPO panel showed suboptimal performance (sensitivity/specificity: Carba5 75.5%/89.0%, CPO panel 78.1%/74.0%). Carba5 test provided specific carbapenemase class assignments but CPO panel failed in 20.3% of cases. Carba5 test and the CPO panel results correlated well with ceftazidime-avibactam minimal inhibitory concentrations (MICs). Concordance for class A/D with MICs was 88.3% for Carba5 and 92.0% for the CPO panel; whereas for class B, it was 86.5% for Carba5 and 76.2% for the CPO panel. In conclusion, FirstAll as the unbiased colony selection impacted carbapenemase testing. With FirstAll, the diagnostic performance of either Carba5 or the CPO panel was compromised. The utilization of ceftazidime-avibactam guided by either the CPO panel or Carba5 was appropriate.

**Importance:** The increasing carbapenemase-producing organisms (CPO) is concerning due to high mortality rates and limited treatment options. Precise testing for CPO is crucial not only for antibiotic treatments but also for infection control. However, discrepant results for an individual overtime or even intra-specimen are found in either phenotypic or genetic testing, posing considerable challenge in clinical management. Based on the colonization-infection model of CPO infections, there would be strain heterogeneity in an individual. On top of the heterogeneity, the single colony selection method in conventional CPO testing would be the source of discrepancy and bias. To test the hypothesis, we proposed FirstAll method as the unbiased colony selection method. We demonstrated that FirstAll corrected around 10% false-negative cases. Lower diagnostic performances of CPO tests were also found in comparison to previous related studies. The study revealed that colony selection would have considerable impacts on CPO testing.

## Introduction

Carbapenemases are β-lactamase enzymes that break down carbapenem antibiotics including penicillins and cephalosporins and belong to molecular class A, B, and D β-lactamases. Carbapenems are effective antibiotics that are used in the treatment of multidrug resistance (1). Carbapenemase-producing (CP) organisms, especially the members in *Enterobacteriaceae* family (CP-CRE) are a global public health concern (1) with its prevalence increasing rapidly (2). Carbapenemase mechanism testing is important for carbapenem-resistant *Enterobacteriaceae* (CRE) prevention because CP-CRE disseminates more readily than non-CP-CRE requiring an intensive infection control approach (3) and has poor prognosis (4). The discrepancies between different carbapenemase tests, such as modified carbapenem inactivation method (mCIM)/EDTA-modified carbapenem inactivation method (eCIM), Carba5, and molecular testing arise from the underlying mechanism (5,6) and from the variations in the diagnostic accuracy. Genetic tests are more sensitive whereas phenotypic tests are only relevant for the functionality (7). Currently, the low degree of automation for the tests contributes to the inter-operator variations (8). Besides, selection bias results in picking colonies from an isolate with heterogeneity leading to discrepancy.

In clinical microbiology laboratories, typically bacteria of the same species are assumed to be homogeneous (9). However, with evolution of many infections, it becomes complex in ageing patients with multiple medical conditions, underlying immunosuppression induced by longer exposure to antibiotics treatment (10). Multiple factors impose strain on bacteria that alter the interaction between microorganisms and hosts. In response to the novel stresses, bacteria evolve variety of strategies to survive *in vivo* (11). This transforms a relatively homogenous bacterial colony to heterogenous that are no longer genetically identical. Theoretically, bacterial heterogeneity is an overlooked reality that is worthy of detailed investigation.

Bacterial heterogeneity has raised considerable attention in recent years. The genetic and phenotypic diversity could be identified between the colonies in carbapenem resistant *Klebsiella pneumoniae* (CRKP) bloodstream infections (BSIs) (12).The heterogeneous strains also exhibited significant differences in virulence in an animal model. The data suggest a new paradigm of CRKP population diversity during BSIs that challenges the single-organism hypothesis and has implications for understanding antibiotic resistance and pathogenesis. Culture-based detection methods are time-consuming, with limited intra-sample abundance, strain diversity information and uncertain sensitivity (13).

Similarly, in a study of enteric vancomycin resistant *Enterococcus* (VRE), clonal diversity and dynamic changes were observed over several weeks (14). These findings suggest that traditional typing methods that analyze one isolate per patient may be insufficient for outbreak surveillance of VRE in highly vulnerable patients. Together, these studies (12–14) highlight the importance of considering within-host microbial diversity and bacterial heterogeneity in the context of infectious disease. They challenge traditional assumptions about the causes and mechanisms of infectious diseases. Moreover, the data suggest that within-host bacterial heterogeneity would be the cause of discrepancy between different lab tests. New approaches to detection and surveillance are necessary. To increase the ability of detecting the complexity of bacterial populations within individual hosts, we propose a novel and non-biased colony collection method (i.e. FirstAll).

Initially, the discrepancy between FirstAll and the conventional colony-picking method was ascertained by comparing FirstAll method with the common carbapenemase tests for six bacterial species. This study comprehensively analyses the performance of various carbapenemase tests and contribute to our understanding of within-host bacterial heterogeneity and its impact on diagnostic methods.

## Materials and Methods

### Strain re-isolation and species identification

Potential seven CPO species - *Klebsiella pneumoniae, Escherichia coli, Enterobacter cloacae complex*, *Klebsiella aerogenes, Klebsiella oxytoca, Citrobacter freundii, Citrobacter koseri*, were re-isolated from bacterial bank. The selection bias was avoided by collecting all the bacterial lawn from agar plates. Isolates were stored at −70°C until analysis. Fresh colonies grown on BBL™ Trypticase™ Soy Agar with 5% Sheep Blood (TSA II) (Becton Dickinson, MD, USA) for 24 hours were picked and smeared onto a MALDI target plate in thin films for identification of bacterial species as described (15).

### Conventional colony and unbiased colony selection methods

We proposed an unbiased bacterial colony-selecting method prior to further analyses. As mentioned above, we have stored the strains with an unbiased method (i.e. scratching all the bacterial lawn off from the agar plates). On this basis, we inoculated the storage onto agar plates and reconfirmed the bacterial species with MALDI-TOF. On the agar plates, furthermore, we adopted two different colony selection methods, including the “conventional method” and the unbiased selection method (referred to as the “FirstAll method” (Figure 1)). While doing the “conventional method,” several single colonies only were collected for further carbapenemase analyses. For the “FirstAll method,” we scratched over the first lawn on agar plates with 10 µl loops. The direction of collecting is vertical to the streaking of the first lawn. Thus, this collection method theoretically maximizes the collection of various strains. The FirstAll method was adopted as the colony selection method for the carbapenemase tests.

**Figure 1.**
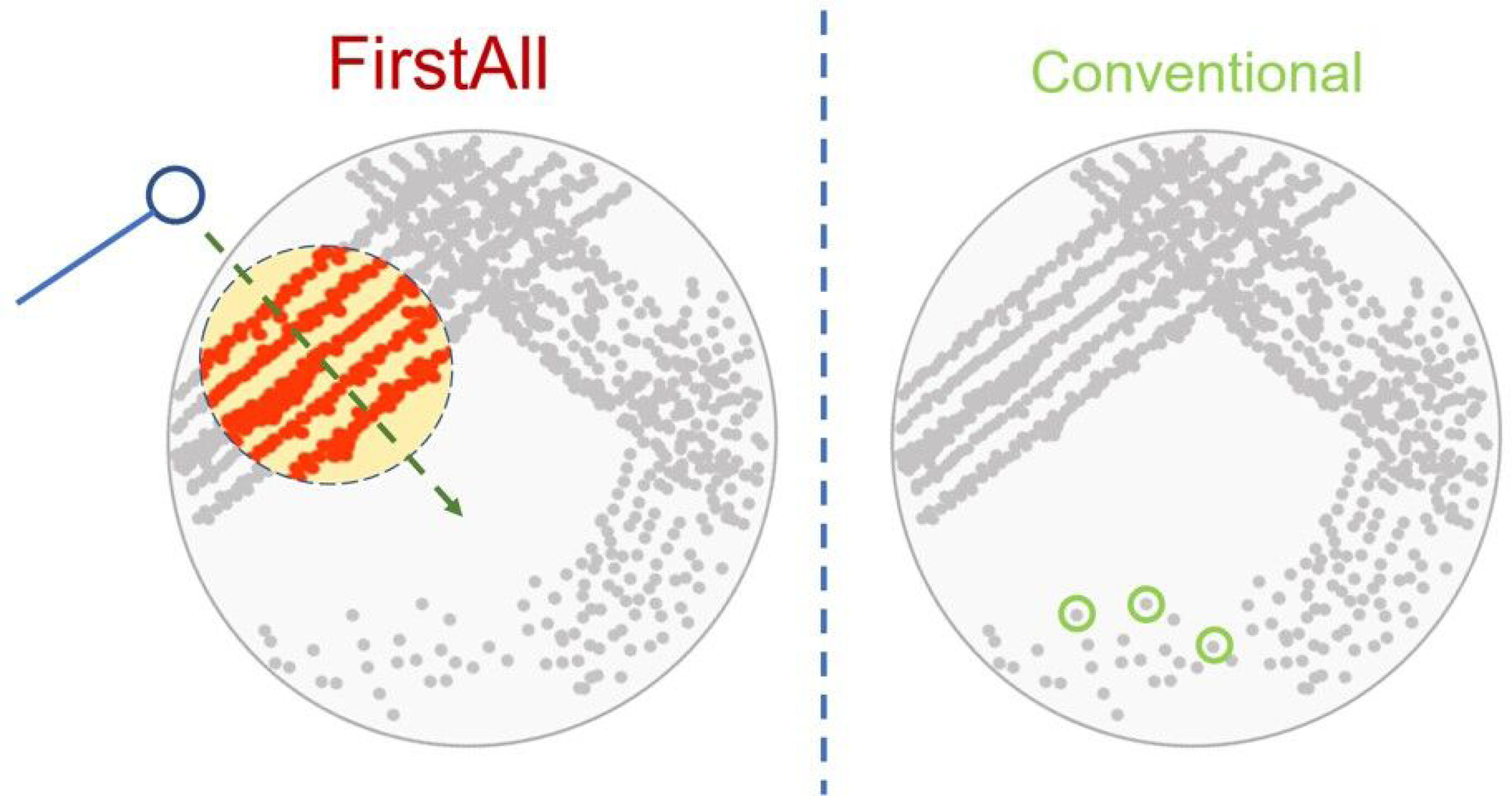
Schematic illustration of FirstAll method for unbiased colony selection. We propose a FirstAll method to increase bacterial diversity and avoid selection bias prior to analyses. In the FirstAll method, we scratch on the agar with a direction vertical to the streaking of the first streak area. By contrast, only several single colonies are picked up for a lab test.

The mCIM and eCIM tests were conducted on bacterial isolates as described by CLSI for *Enterobacterales* (5). The test was performed in triplicate and the results were interpreted by three independent technicians.

### BD CPO panel

The BD Phoenix CPO panel (BD, NJ, USA) includes the ID/NMIC 504 test for detecting carbapenemase activity, and the NMIC 500 test for Ambler classification. The CPO panel utilizes meropenem, doripenem, temocillin and cloxacillin, alone and in combination with various chelators and beta-lactamase inhibitors for the detection and classification of CPOs. Specific Ambler classification was not provided for the cases with negative results. The test results were interpreted using BD EpiCenter™ software (BD) as per the latest CLSI or EUCAST guidelines.

### Carba5 test

For the CARBA-5 test (NG-Biotech, Guipry, France) five drops of the extraction buffer was mixed with a full 1-µl inoculation loop of CPO bacteria culture, and from this 150 µl was transferred into the CARBA-5 cassette. The results were read after 15 minutes of incubation at room temperature.

### Multiplex PCR (M-PCR)

A multiplex PCR targeting important carbapenemase genes was designed (Table 1). DNA was extracted from isolated colonies with a QIAamp DNA minikit (Qiagen, Taiwan. The concentration and purity of the extracted DNA were verified using a Nanodrop-ND1000 (Thermo Fisher Scientific, Waltham, MA, USA). PCR assays were be performed with 100 ng of genomic bacterial DNA, 10 pmol of each primer (Mission Biotech), 10 μl of Phusion PCR Master Mix (Thermo Fisher) and ultra-pure water to a final reaction volume of 20 μl. The PCR conditions included an initial denaturation at 98 °C for 1 minute followed by 30 cycles of 98 °C for 1 second, 58 °C for 5 seconds, and 72 °C for 10 seconds, with a final extension at 72 °C for 1 minute. PCR amplicons were electrophoresed on a 1.5% (w/v) agarose gel in the TAE buffer. A 100 bp DNA ladder (Promega Corporation, Madison, USA) was included in each run. After electrophoresis, the gels were photographed under UV at 260 nm.

**Table 1:**
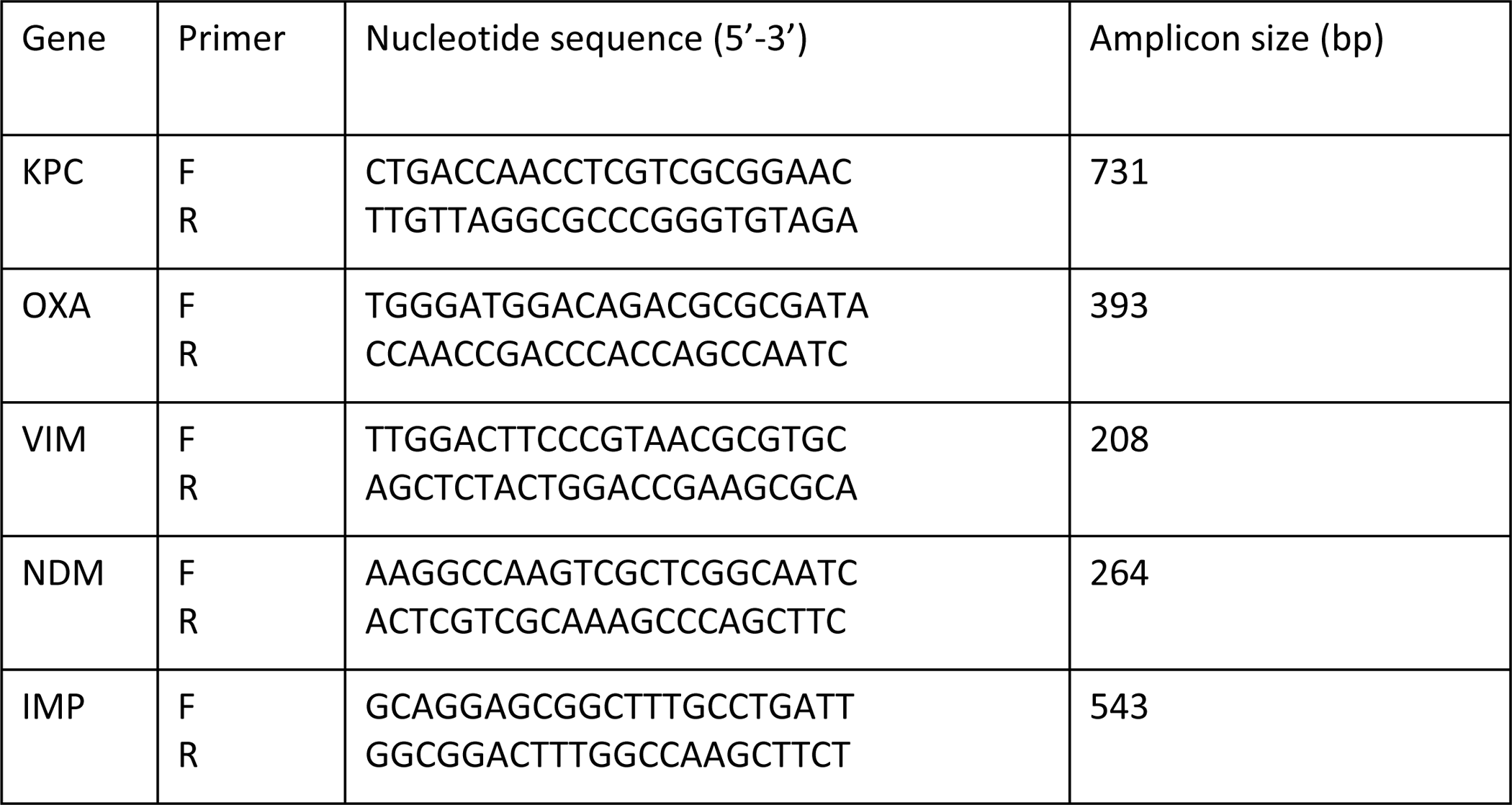
Primers sequence for top 5 carbapenemase genes. We designed primer sets for the five most common carbapenemase genes (i.e., KPC, OXA, VIM, NDM, IMP). The size of the expected amplicons was designed to be different so that the amplicons could be distinguishable in a multiplex PCR setting. F: forward; R: reverse.

### MIC of antibiotics

The minimum inhibitory concentration (MIC) was tested with BD Phoenix CPO panel NMIC 500 (BD) for 25 different antibiotics, including ceftazidime-avibactam and carbapenems, using true doubling dilutions. CPO with class A/D carbapenemase is typically susceptible to ceftazidime-avibactam, whereas class B carbapenemase is resistant to ceftazidime-avibactam. Cutoff of ≤8/4 μg/ml was used to call susceptible to ceftazidime-avibactam, while ≥16/4 μg/ml was interpreted as resistant to ceftazidime-avibactam.

## Results

### Discrepancy of mCIM results based on different colony selection methods (FirstAll method versus Conventional method)

The *K. pneumoniae* isolates were used as the CPO model to evaluate the idea of unbiased colony selection (Figure 2). The mCIM results from FirstAll concurred with conventional colony selection methods (88.43%, 260/294). Discrepancy was noted in 34 out of 294 isolates. Amid the discrepant cases, 31 (10.54% (31/294)) isolates showed mCIM (+) with FirstAll, but were reported mCIM (−) by conventional method. The results indicate that 10.54% *K. pneumoniae* isolates tested falsely as negative for carbapenemase by the conventional colony picking-up method.

**Figure 2.**
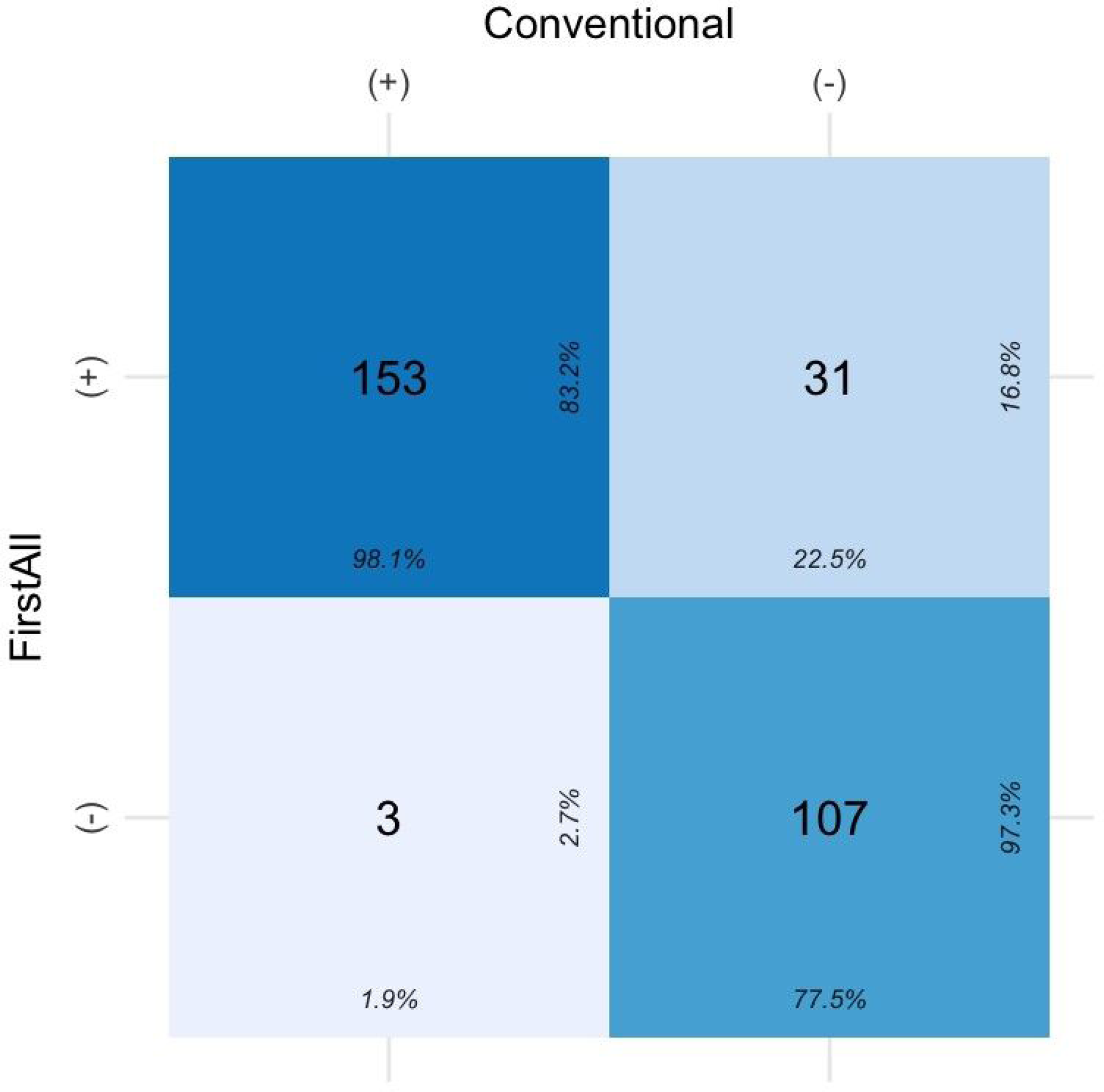
Discrepancy of mCIM results between FirstAll and conventional colony selection methods. Most of the *K. pneumoniae* isolates (both methods show positive: 153; both methods show negative: 107) show concordant mCIM results no matter whether FirstAll or conventional colony selection method is used. Of note, 31 out of 294 isolates reveal mCIM positive for the FirstAll method but negative for the conventional method, indicating FirstAll would be an unbiased colony selection method.

### Carbapenemase testing comparison (CPO panel versus mCIM/eCIM)

For most CPO species except K. pneumoniae, the CPO panel agreed with mCIM/eCIM well (Figure 3). The CPO panel also provided exact classifications (i.e., Class A or D, Class B) for most CPO species. In case of mCIM(+)/eCIM(−) *K. pneumoniae* isolates, nearly half the isolates (43/88) were from class A or D. However, for the other half of *K. pneumoniae* isolates with mCIM(+)/eCIM(−) the class were not determined. Similarly, in *K. pneumoniae* isolates with mCIM(+)/eCIM(+), 38.60% (22/57) were class A or D while only 13.56% (8/57) were class B. Of note, for 47.37% (27/57) *K. pneumoniae* isolates with mCIM(+)/eCIM(+), carbapenemase was detected by the CPO panel but the class was not determined.

**Figure 3.**
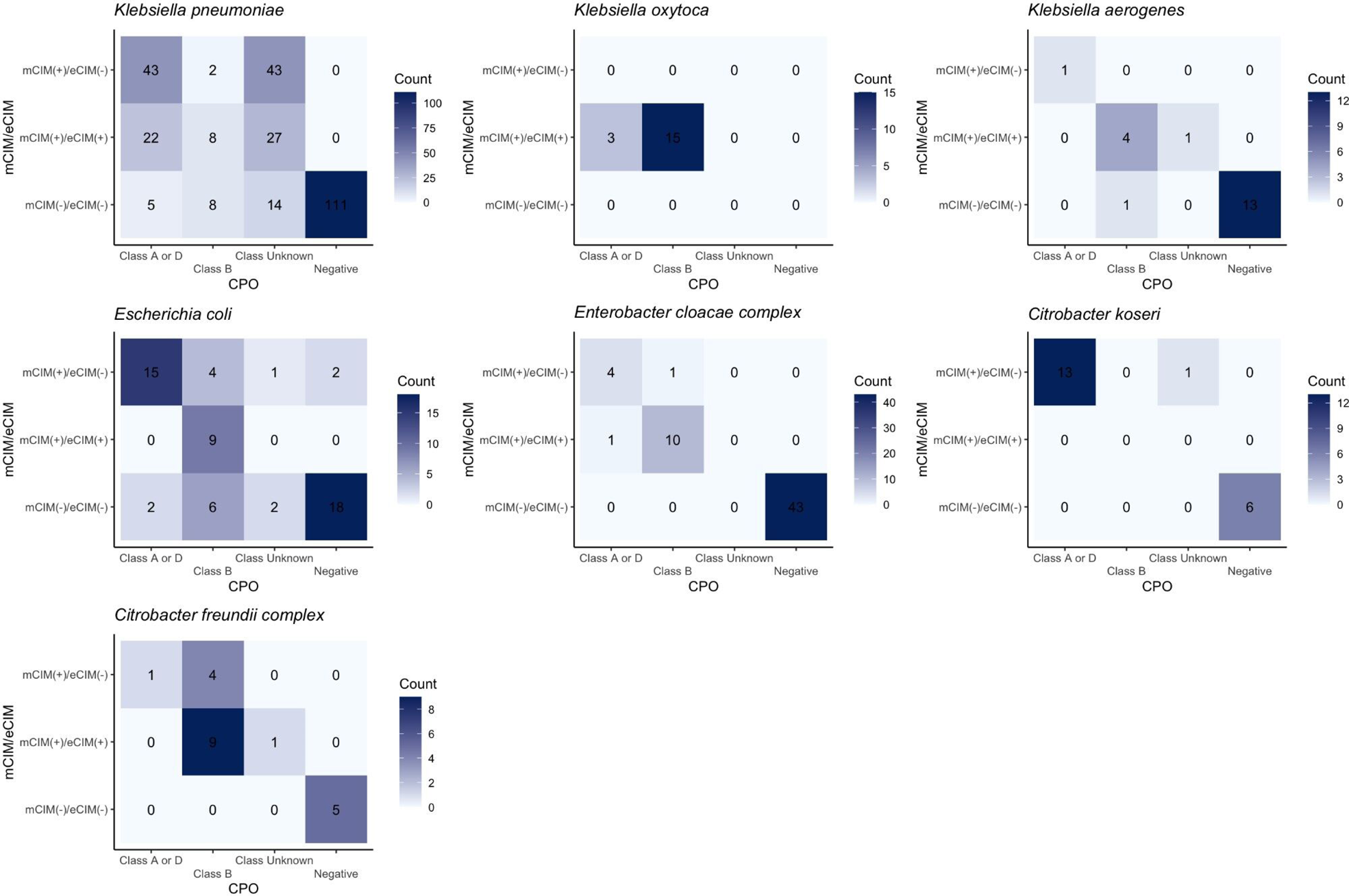
Comparison between CPO panel and mCIM/eCIM. General categorical agreements between the CPO panel and mCIM/eCIM can be found along the diagonal lines except in *K. pneumoniae*. The CPO panel can detect the existence of carbapenemase in *K. pneumoniae*, but the class cannot be determined. Specifically, around half of *K. pneumoniae* isolates (43/88) with mCIM(+)/eCIM(−) (i.e., regarded as class A/D carbapenemase) are categorized as “Class Unknown”; twenty-seven out of 57 *K. pneumoniae* isolates with mCIM(+)/eCIM(+) (i.e., regarded as class B carbapenemase) are categorized as “Class Unknown.”

### Carbapenemase testing comparison (CPO panel versus M-PCR)

The phenotypic results of the CPO panel were also compared with the genotypic results by using PCR as the reference method. The comparisons in different species were joined together to avoid scattered results (Figure 4A). For the “Class A or D” called by the CPO panel, 72.32% (81/112) was in line with MPCR results, while 2.68% (3/112) was called IMP/NDM/VIM, 18.75% (21/112) was categorized as “Others” (i.e. multiple genes detected), and 6.25% (7/112) was called negative. For the “Class B” category, 60% (48/80) agreed with MPCR, while notably, 18.75% (15/80) were called negative. With the FirstAll method adopted, a considerable proportion (20.33%, 100/492) was called positive for carbapenemase by the CPO panel but failed to be specifically categorized (i.e., the “Class Unknown”).In cases defined as negative by the CPO panel, the majority (65.50%, 131/200) was also negative by MPCR. However, still 34.5% of isolates were found harboring carbapenemase genes.

**Figure 4A.**
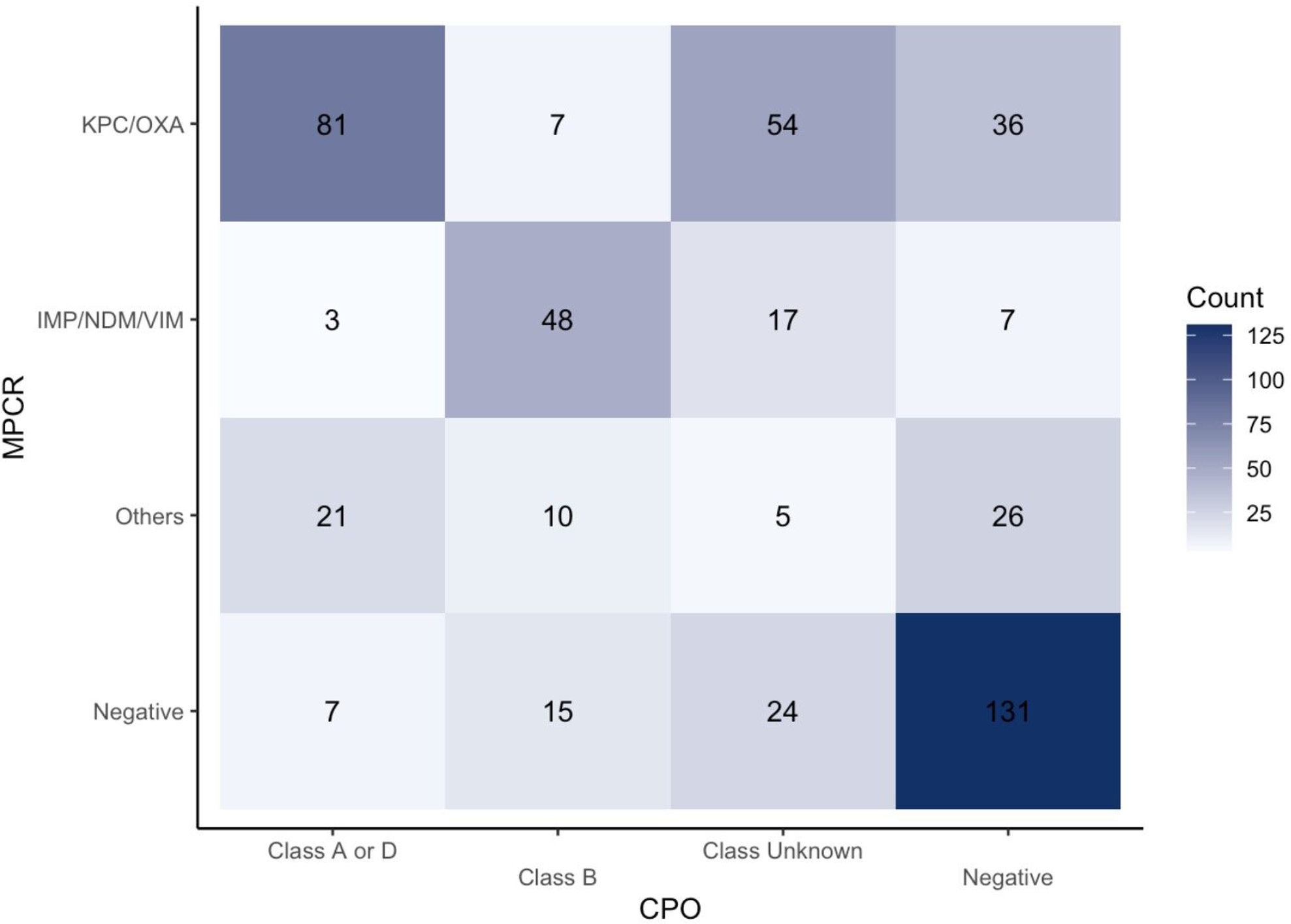
Comparison between CPO panel and MPCR. The CPO panel and MPCR agreement are generally fair except for the “Class Unknown” of the CPO panel. The “Class Unknown” accounts for 20.33% [100/492] of the all isolates. The overall sensitivity and specificity are 78.10% [246/315] and 74.01% [131/177], respectively.

### Carbapenemase testing comparison (Carba5 versus M-PCR)

The performance of Carba5 was evaluated by using MPCR as the reference method. A concordant rate of around 73% was noted for all the categories (Figure 4B). Specifically, 79.75% (130/163) of cases where the Carba5 test showed a KPC/OXA result also showed the same result on the MPCR. For IMP/NDM/VIM class, 82.89% (63/76) of IMP/NDM/VIM cases tested by Carba5 revealed the same pattern on MPCR. For all the cases called negative for carbapenemase by Carba5, 67.36% (161/239) cases also showed negative results in MPCR. Of note, around 10% of cases that were determined as a specific class by Carba5 were classified as Others by MPCR (for class KPC/OXA: 16/163; for class IMP/NDM/VIM: 9/76), indicating more multiple carbapenemase classes detected by MPCR.

**Figure 4B.**
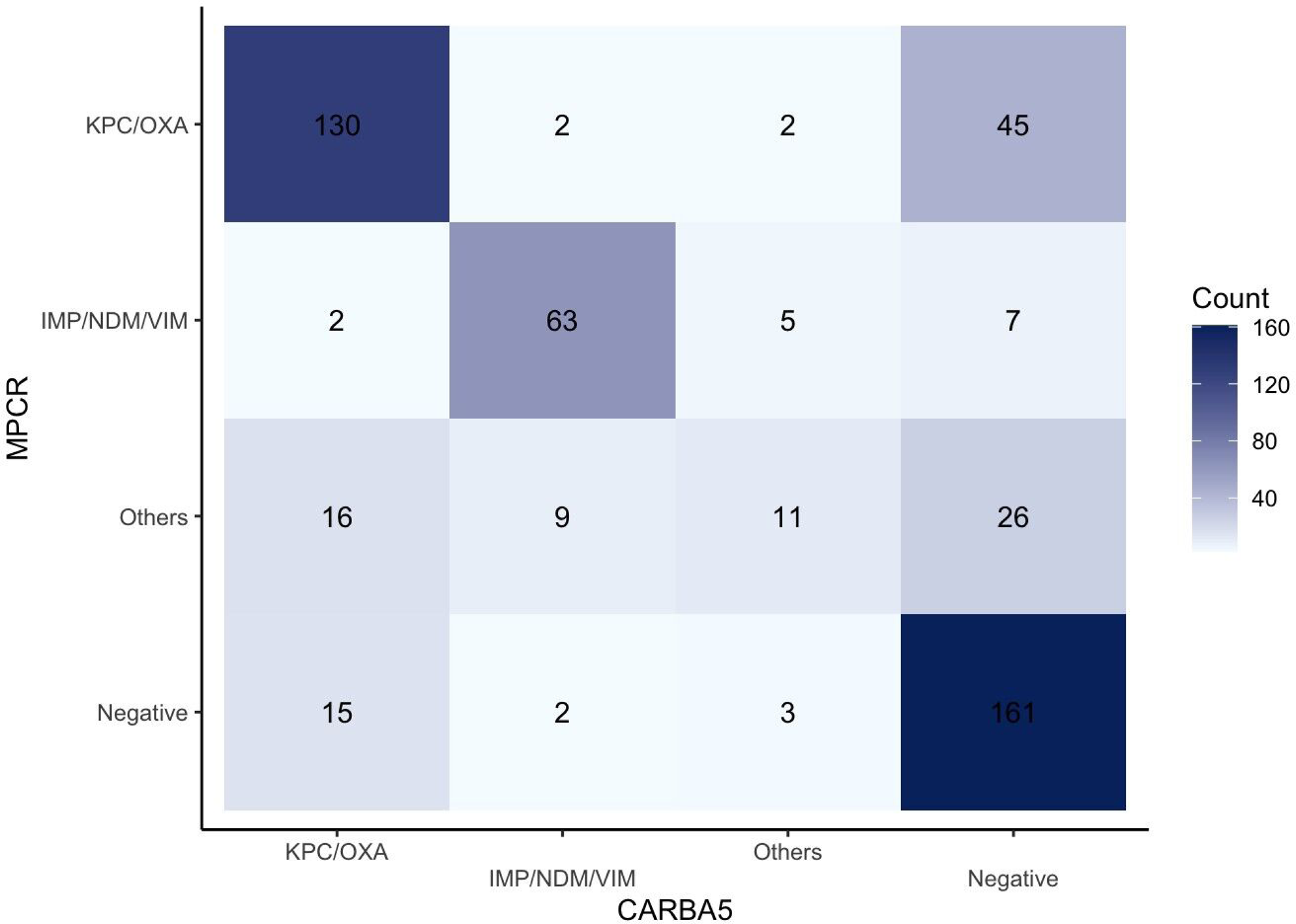
Comparison between Carba5 and MPCR. The agreement between Carba5 and MPCR is high for all the categories. The overall sensitivity and specificity are 75.47% [240/318] and 88.95% [161/181], respectively.

### Carbapenemase testing comparison (CPO panel versus Carba5)

The CPO panel results generally agreed with Carba5 results (Figure 5A) except for the “Class Unknown” category. Specifically, 85.71% (96/112) “Class A or D,” 66.25% (53/80) “Class B,” and 95.00% (190/200) “Negative” results called by the CPO panel were also in line with the results by using Carba5. Notably, the agreement between the two methods was especially low for class B carbapenemase. Moreover, a considerable proportion (19.15%, 45/235) of Carba5 negative isolates were called positive by the CPO panel. Further comparison was done for the MPCR-positive isolates (Figure 5B). For the isolates that were considered carbapenemase positive by MPCR (the reference method), agreements between the CPO panel and Carba5 in “Class A or D” and “Class B” were 88.57% (93/105) and 78.46% (51/65), respectively. Generally, the agreements between the CPO panel and Carba5 were similar in either the MPCR-positive subgroup or in the whole population (70.73% (348/492) and 69.46% (210/315), respectively). Of note, still 19.05% (60/315) of MPCR-positive cases were not detected by the CPO panel or Carba5.

**Figure 5A.**
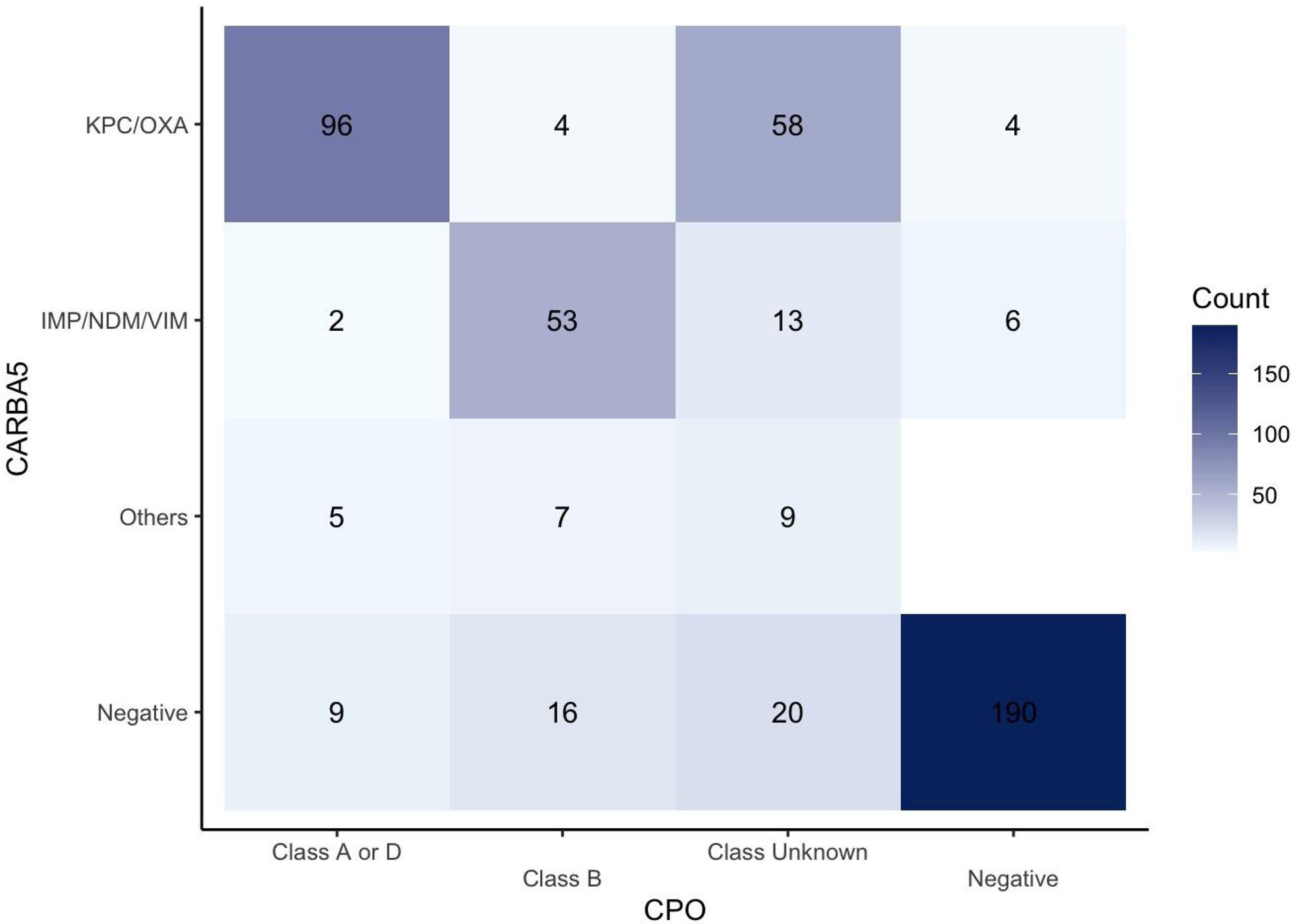
Comparison between CPO panel and Carba5. The agreement between the two methods is generally good for the “Class A or D” and “Negative” categories, but the agreement is low for the “Class B” category. Moreover, for the isolates called positive by the CPO panel, 34.25% (100/292) isolates are called positive for carbapenemase but class unknown.

**Figure 5B.**
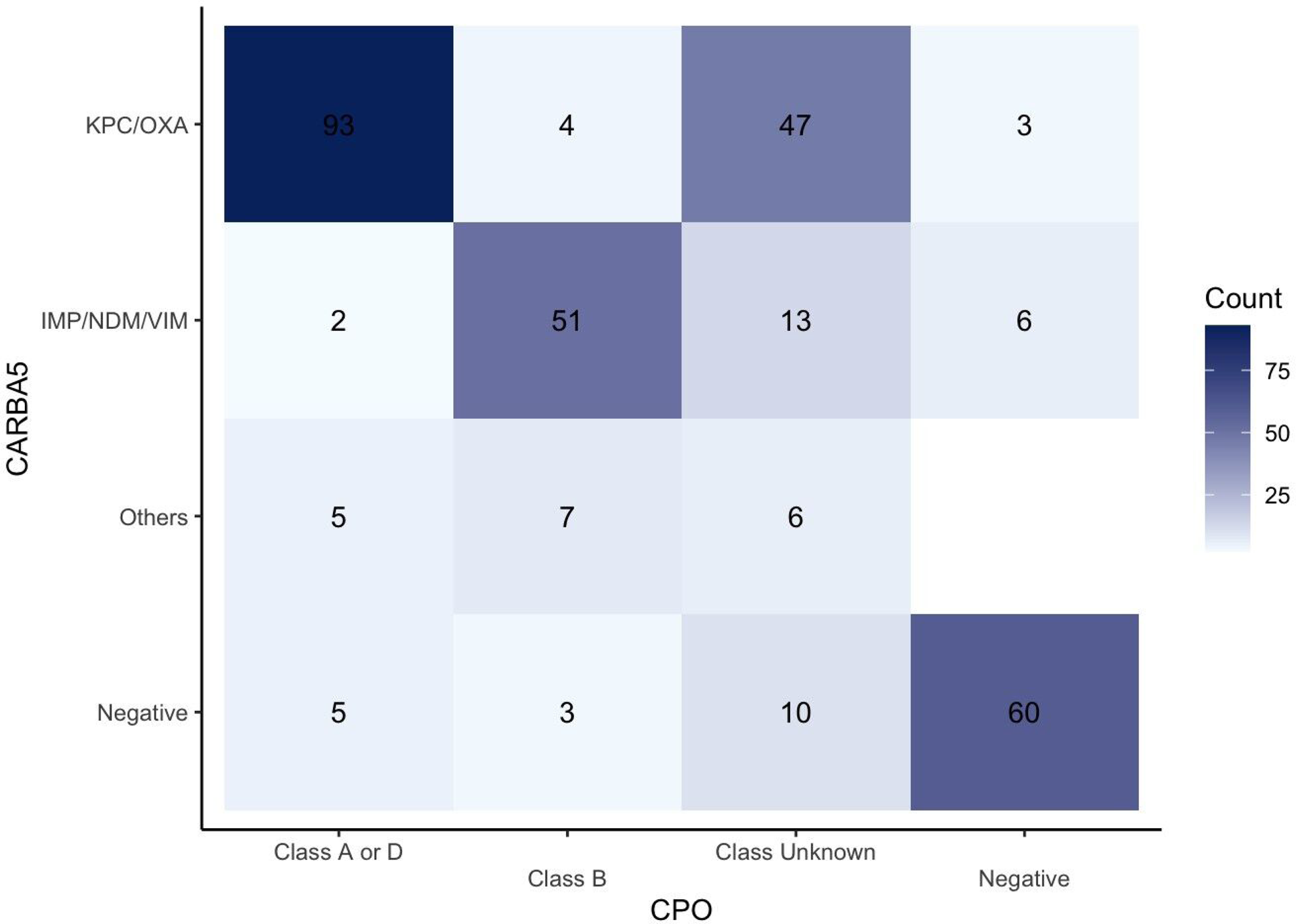
Comparison between CPO panel and Carba5 on the MPCR positive strains.

### Association between CPO panel results and MICs of Ceftazidime-avibactam

Ceftazidime-avibactam can be used to treat CPOs with class A/D carbapenemase but not for class B carbapenemase. Thus, an agreement between the carbapenemase test and ceftazidime-avibactam MIC is important for clinical decisions. For “Class A or D,” called by the CPO panel, 92.04% (104/113) cases were susceptible to ceftazidime-avibactam (Figure 6A). For “Class B,” called by the CPO panel, 76.19% (64/84) cases were resistant to ceftazidime-avibactam, and still 23.81% (20/84) cases were interpreted as susceptible. Of note, for the “Class Unknown’’ category, a biurnal MIC distribution was found: 61.54% (64/104) cases were susceptible to ceftazidime-avibactam, while still 26.92% (28/104) strains were resistant to ceftazidime-avibactam. A similar pattern of agreement can be observed for Carba5 and ceftazidime-avibactam MIC (Figure 6B). On top of the pattern, Carba5 also showed high agreement with ceftazidime-avibactam MIC: 88.27% (143/162) “KPC/OXA” cases were susceptible to ceftazidime-avibactam, while 86.49% (64/74) “IMP/NDM/VIM” cases were resistant to ceftazidime-avibactam. For the “Others” category, multiple carbapenemase classes were identified, and mostly (61.90% (13/21)) presented resistant to ceftazidime-avibactam.

**Figure 6A.**
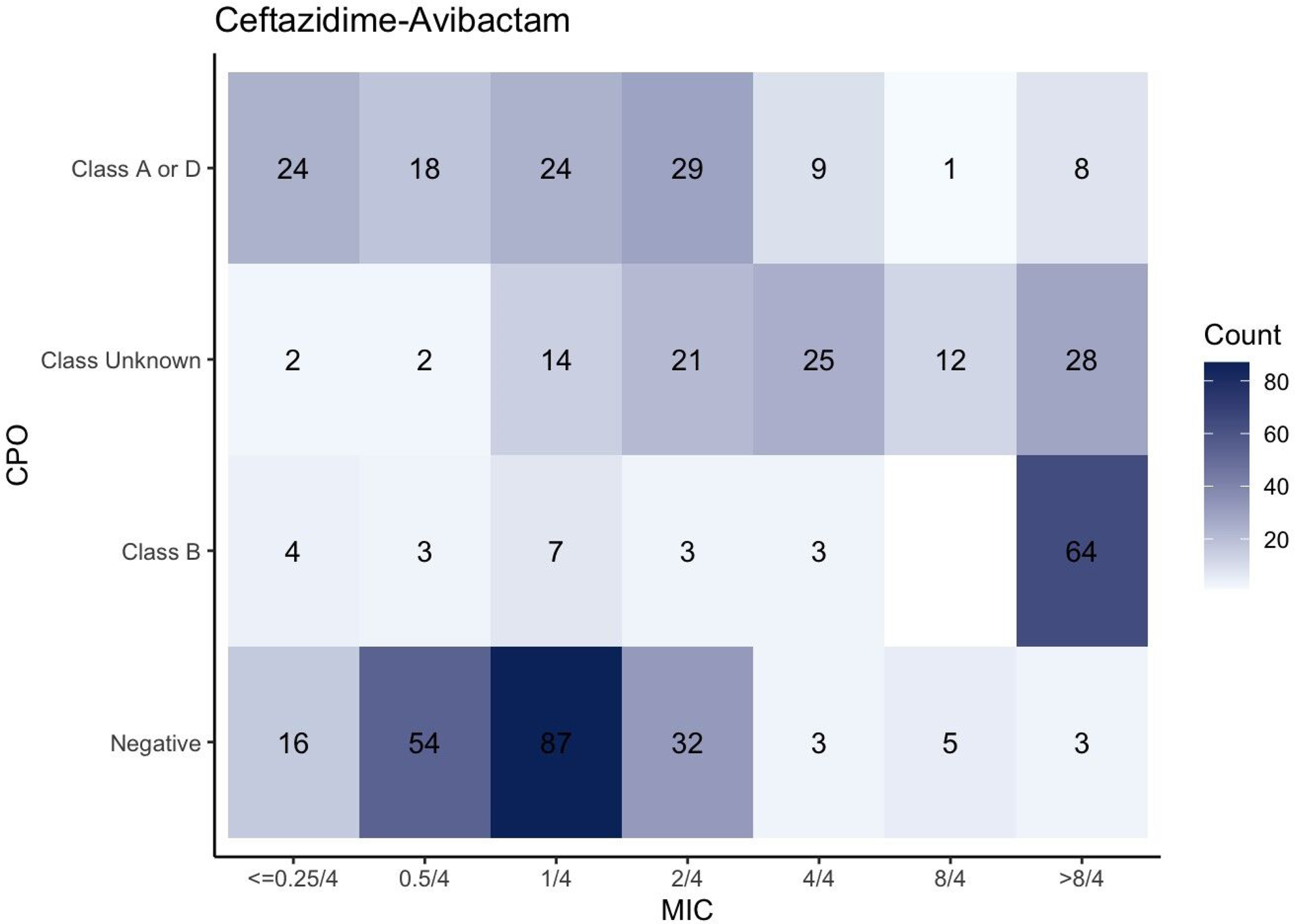
MICs of ceftazidime-avibactam in different classes of CPO panel.

**Figure 6B.**
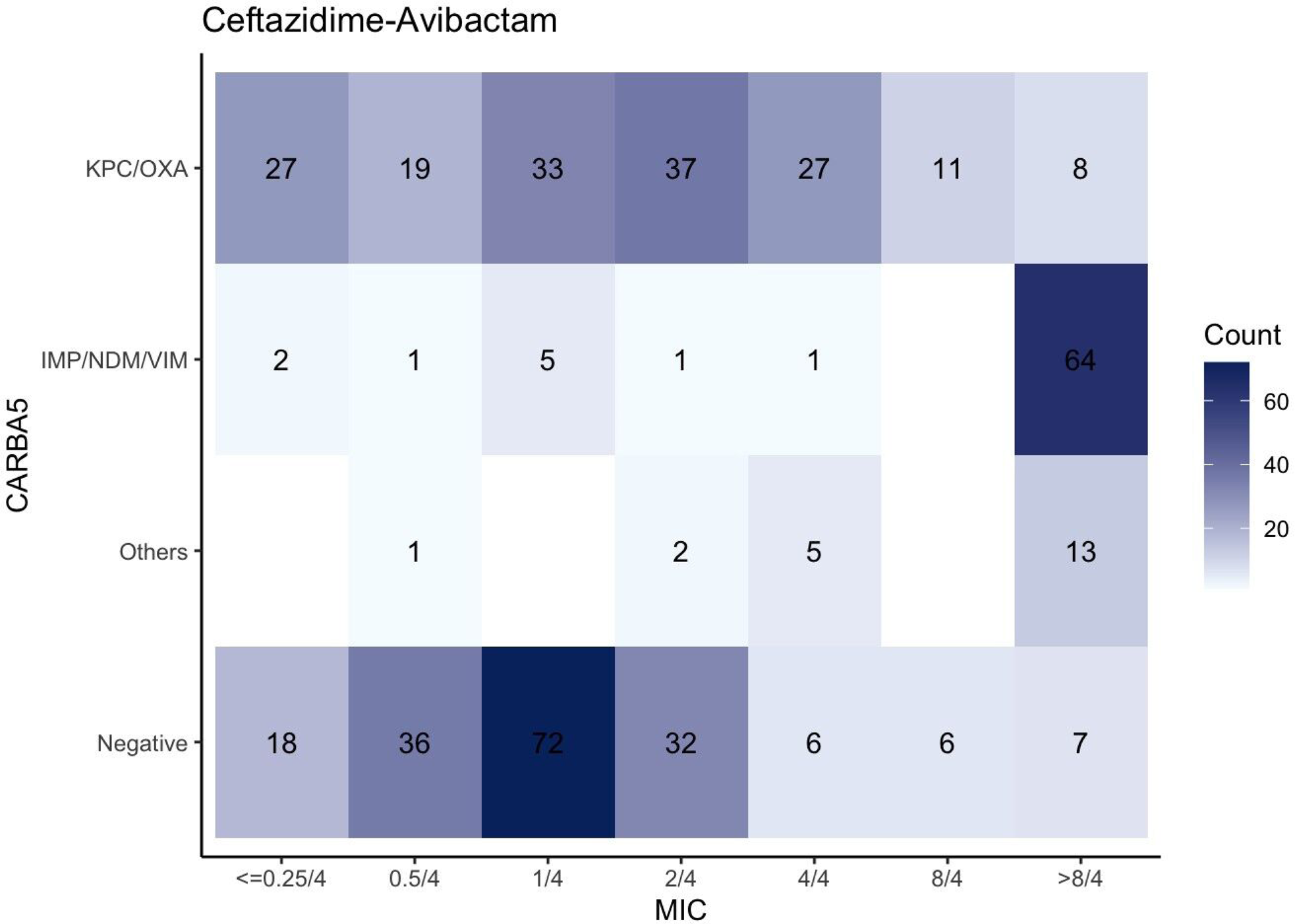
MICs of ceftazidime-avibactam in different classes of Carba5.

## Discussion

CPOs pose a significant threat to public health due to their ability to break down carbapenem antibiotics, which are considered as the last line of defense against serious infections. In this study, we compared multiple carbapenemase tests based on an unbiased colony selection method (i.e. FirstAll) and investigated the impact of within-host bacterial heterogeneity on diagnostic methods of carbapenemase. Our findings shed light on the impact and importance of the unbiased colony selection method and as well provided insights into the complexity of bacterial populations within individual hosts.

Colonies on an agar plate with the same morphology are considered identical, a fundamental assumption in clinical microbiology laboratories, that has been challenged in recent years. Blood stream infections of CRKP (CRKP-BSI) does not meet this criteria (12). Gastrointestinal colonization with *K. pneumoniae* is a major predisposing risk factor for infection and a hub for the dispersal of resistance (13) and are considered endogenous rather than exogenous. The KP strains colonize the host’ gut for a long time (13). The KP strains typically harbor multiple and diverse antibiotic resistance mechanisms for survival against antibiotics challenges for multiple times (13). The KP strains reach a colonization balance and no survival advantage for one KP strain than another. Once the host’ immunity is weakened, the diverse KP strains will disseminate out from the gut and get into the bloodstream. In the case, there is considerable strain diversity within a specimen, even the morphologies of the strains are identical.

To avoid selection bias in lab tests, we developed FirstAll to increase the strain diversity on colony selection. The results showed that around 10% (31/294) KP strains were called positive in FirstAll but negative in the conventional method. The 10% discordance between the colony selection methods indicates that the conventional colony selection method is insufficient for detection of intra-specimen KP strain diversity. The inability of detecting diverse strains leads to incorrect antibiotic selection and unfavorable clinical outcomes. Our findings posed that FirstAll would be a more comprehensive, unbiased, and adequate colony selection method. By contrast, the utility of the conventional colony selection method needs to be reinvestigated to avoid false negatives of carbapenemase detection.

Intra-specimen strain diversity would account for the discrepancy between carbapenemase tests. In the comparison study between CPO panel and mCIM/eCIM, we showed the subgroup comparisons for all the bacterial species. Generally, the CPO panel results agreed well with the mCIM/eCIM results. Interestingly, KP was the only bacterial species that the CPO panel did not agree well with mCIM/eCIM, highlighting the species-specific nature of these tests. The CPO panel did not allocate specific classes for around 50% KP species. The high chance of being called “Class Unknown” was not noted in other species. Tracing back to the working mechanism of the CPO panel, cases with carbapenemase positive but class unable to be determined are categorized as “Class Unknown”. The high percentage of “Class Unknown” in KP strains implies that there were diverse carbapenemase classes by using FirstAll as the colony selection method. While other bacterial species did not show a high percentage of “Class Unknown”, the underlying mechanism between the bacterial species is worthy of further investigation.

With M-PCR as the reference method, both CPO panel and Carba5 showed high concurrence with M-PCR when specific classes were allocated. The major difference between CPO panel and Carba5 arises from strains, as multiple carbapenemases classified by Carba5 are classified as “Class Unknown” by the CPO panel. The information of specific classes is beneficial in choosing antibiotic (e.g. ceftazidime-avibactam). For the M-PCR positive subgroup, 19.05% cases were neither reported in CPO panel nor in Carba5. Adopting FirstAll colony selection method, the diagnostic performance of the CPO panel or Carba5 was lower than those reported in previous studies (16,17). The hypothesis is that while FirstAll unbiasedly collects more colonies than the conventional colony collection method, in some cases the minor variants account for only a small portion. The minority variants may be detected by high sensitive methods (such as M-PCR) but not by phenotypic assays. The clinical impacts of the extra carbapenemase detection due to the unbiased colony selection would be an interesting topic to investigate.

Several limitations should be addressed. First, the CPO strains were isolated from a single referral medical center in East Asia. Given the fact that local circulating strains may differ considerably between areas, hospital type and local epidemiology should be taken into consideration in interpreting the results. Second, genomic information was not tested, especially for the cases showing multiple carbapenemases. It is not clear whether the multiple carbapenemases are derived from a single strain or from multiple strains that were collected by FirstAll method. Genomic study, preferably long-read sequencing would be necessary to reveal the genetic context between the carbapenemase genes. Our study would be the first to reveal the impact of colony selection method on carbapenemase tests. Besides, the findings emphasize the complexity and potential clinical impacts of the carbapenemase tests.

## Conclusion

The study highlights the importance of unbiased colony selection methods in carbapenemase testing. Under an unbiased colony selection method, diagnostic performance of either Carba5 or the CPO panel was compromised. Class-specific detection by Carba5 would aid clinical decision making. Ceftazidime-avibactam use based on either the CPO panel or Carba5 is suitable.

## Supplement

**Figure 1. Simulated and actual gel electrophoresis for the top 5 carbapenemase amplicons.** We simulate the positions for the amplicons (left) and demonstrate the results (right) of single plex PCR for the top 5 carbapenemase genes. The positions are concordant with the simulation. Based on the single-plex results, we design a multiplex PCR.

## Reference

1. Bonomo RA, Burd EM, Conly J, Limbago BM, Poirel L, Segre JA, et al. Carbapenemase-Producing Organisms: A Global Scourge. Clin. Infect. Dis. Off. Publ. Infect. Dis. Soc. Am. 66, 1290–1297 (2018).

2. Cui, X., Zhang, H. & Du, H. Carbapenemases in Enterobacteriaceae: Detection and Antimicrobial Therapy. Front. Microbiol. 10, 1823 (2019).

3. CRE Technical Information | CRE | HAI | CDC. https://www.cdc.gov/hai/organisms/cre/technical-info.html (2021).

4. Tamma PD, Goodman KE, Harris AD, Tekle T, Roberts A, Taiwo A, et al. Comparing the Outcomes of Patients With Carbapenemase-Producing and Non-Carbapenemase-Producing Carbapenem-Resistant Enterobacteriaceae Bacteremia. Clin. Infect. Dis. Off. Publ. Infect. Dis. Soc. Am. 64, 257–264 (2017).

5. CLSI. Performance Standards for Antimicrobial Susceptibility Testing. 33th ed. CLSI supplement M100. Wayne PA Clin. Lab. Stand. Inst. (2023).

6. Maurer, F. P., Castelberg, C., Quiblier, C., Bloemberg, G. V. & Hombach, M. Evaluation of Carbapenemase Screening and Confirmation Tests with Enterobacteriaceae and Development of a Practical Diagnostic Algorithm. J. Clin. Microbiol. 53, 95–104 (2015).

7. Kalpana S, Lin WY, Wang YC, Fu Y, Lakshmi A, Wang HY. Antibiotic Resistance Diagnosis in ESKAPE Pathogens—A Review on Proteomic Perspective. Diagnostics 13, 1014 (2023).

8. Lippi, G. & Rin, G. D. Advantages and limitations of total laboratory automation: a personal overview: Clin. Chem. Lab. Med. CCLM 57, 802–811 (2019).

9. Kragh KN, Alhede M, Rybtke M, Stavnsberg C, Jensen PØ, Tolker-Nielsen T, et al. The Inoculation Method Could Impact the Outcome of Microbiological Experiments. Appl. Environ. Microbiol. 84, e02264–17 (2018).

10. Bloom, D. E. & Cadarette, D. Infectious Disease Threats in the Twenty-First Century: Strengthening the Global Response. Front. Immunol. 10, 549 (2019).

11. Mwangi MM, Wu SW, Zhou Y, Sieradzki K, de Lencastre H, Richardson P, et al. Tracking the in vivo evolution of multidrug resistance in Staphylococcus aureus by whole-genome sequencing. Proc. Natl. Acad. Sci. U. S. A. 104, 9451–9456 (2007).

12. Cheng S, Fleres G, Chen L, Liu G, Hao B, Newbrough A, et al. Within-Host Genotypic and Phenotypic Diversity of Contemporaneous Carbapenem-Resistant Klebsiella pneumoniae from Blood Cultures of Patients with Bacteremia. mBio 13, e0290622 (2022).

13. Lindstedt K, Buczek D, Pedersen T, Hjerde E, Raffelsberger N, Suzuki Y, et al. Detection of Klebsiella pneumoniae human gut carriage: a comparison of culture, qPCR, and whole metagenomic sequencing methods. Gut Microbes 14, 2118500 (2022).

14. Both A, Kruse F, Mirwald N, Franke G, Christner M, Huang J, et al. Population dynamics in colonizing vancomycin-resistant Enterococcus faecium isolated from immunosuppressed patients. J. Glob. Antimicrob. Resist. 28, 267–273 (2022).

15. Wang HY, Li WC, Huang KY, Chung CR, Horng JT, Hsu JF, et al. Rapid classification of group B Streptococcus serotypes based on matrix-assisted laser desorption ionization-time of flight mass spectrometry and machine learning techniques. BMC Bioinformatics 20, 703 (2019).

16. Baeza LL, Pfennigwerth N, Greissl C, Göttig S, Saleh A, Stelzer Y, et al. Comparison of five methods for detection of carbapenemases in Enterobacterales with proposal of a new algorithm. Clin. Microbiol. Infect. 25, 1286.e9–1286.e15 (2019).

17. Khalifa HO, Okanda T, Abd El-Hafeez AA, El Latif AA, Habib AGK, Yano H, et al. Comparative Evaluation of Five Assays for Detection of Carbapenemases with a Proposed Scheme for Their Precise Application. J. Mol. Diagn. 22, 1129–1138 (2020).

